# Tetrasulfide Bridge (TTSB) Formation: an Extraordinary Intermediate Structure in the Architecture and Dynamics of the HIV-1’s gp120 molecule

**DOI:** 10.1101/2022.03.14.484205

**Authors:** Harry F. Crevecoeur

## Abstract

Gp120, one of the molecules that execute HIV viral entry in CD4+ cells has a number of allosteric disulfide bridges. In this study we explore the dynamics of these potential disulfide (disulphide) bridges in the 3-D configuration (based on crystallography) and UNSW DBA analyses of various gp120 crystals. The data reveal the existence of a tetrasulfide (tetrasulphide) bridge (TTSB) which, together with a disulfide bridge, keeps two perpendicular beta sheets (namely V3 and V4) approximated in the architecture of the gp120 while allowing safe transfer of energy between the allosteric bonds. Analyses of multiple gp120 crystals reveal the existence of the TTSB to be more as an intermediate as opposed to a constant landmark, which implies more complex functions than just structural attributes. This TTSB, which is observed in various crystals of gp120, in various strains and clades of HIV-1, is demonstrated by various rendering software and some are also detected, reported and characterized by the UNSW Disulfide Bond Analysis (DBA) engine. This tetrasulfide bridge connects the allosteric bond Cys296–Cys331 to Cys385-Cys418, sometimes as CYS331:SG – CYS385:SG, and sometimes as CYS331:SG – CYS418:SG. Moreover in crystals 3TIH, 4LSR and 4R4N, we also observed an intermediate state of the TTSB where it presents a triangular formation in which the sulfur atom CYS331:SG binds to both CYS385:SG and CYS418:SG simultaneously, which represents an important intermediate in the functional dynamics of the gp120 molecule. The presence of this extraordinary structure (TTSB) does point to some intriguing insights in the multifunctional design and mechanics as well as some complex fragilities of the gp120 molecule, exposing a new target for antiviral therapy.

## Introduction

HIV-1, the virus that causes AIDS, enters the human cells primarily through interactions of its surface gp120 with the CD4 receptor as well as the CC-CKR5 receptors (Dragic 1996) [1]. The mechanisms of these interactions have yet to be elucidated. It is known that the binding to CD4 requires and engenders additional conformational changes in the gp120 molecule. According to Kwong et al. who solved the first X-ray crystal structure of the gp120, the structure of the gp120 has no precedent [2,3,4]. In addition the gp120 has some unique features that are not found in any other known protein structure. The bistable flip/switch phenomenon, for example, represents a truly unique feature of the gp120 of HIV-1. The bistable flip/switch phenomenon, described by Reed et al., pertains to a segment of the gp120 molecule (encompassing L416 through W427 *in Kwong’s model – HXBc2 strain of HIV-1*) that can alternate between alpha helix and beta configurations without intermediates [5]. The LPCR motif with L416-P417-C418-R/K419 was deemed an important factor controlling these conformational changes [5,6,7]. The fact that “switch inhibitors”, - which are molecules designed to stabilize this segment in one configuration, thereby preventing the conformational change of the bistable flip/switch, - are shown to also inhibit CD4-binding (von Stosch & Reed 1994, ’96, ’99), strongly supports the possibility that the bistable switch phenomenon is important for CD4-binding [6,7,8]. Moreover a number of residues located in this dynamic segment (C416-W427) are also proven necessary for HIV-1 to bind to CC-CKR5 and CD4 namely Arg419/Lys419, Lys421, Gln422 and Trp427 [9,10]

The interaction of the gp120 with the CD4 and CC-CKR5 receptors is a critical and necessary step for HIV infection of human cells. Allosteric disulfide bonds are involved in these mechanisms [11]. In this study we explore the potential disulfide bridges: Cys378-Cys445, Cys296-Cys331 and Cys385-Cys418) of the HIV gp120 in their 3-D configurations (based on crystallography) and UNSW DBA analysis of the gp120 crystals.

### The Nature of Disulfide Bridges

Disulfide bridges provide vital links between strands in protein architecture. They are formed by oxydoreductase enzymes during protein maturation. Disulfide bridges are expected in protein structures. All disulfide bridges are not equal: there are 20 possible configurations divided in 3 basic types: Spirals, Hooks and Staples [11]. The chi (X angles) between the carbon and sulfur atoms in the joining cysteine residues define the geometry of the disulfide bond. The chi angles are used to estimate the dihedral strain energy (DSE) of the bond. While DSE calculations have been shown to be useful measure of the disulfide strain, they vary widely in different crystallization methods: the mean DSE in disulfides resolved by NMR is twice that of disulfides in crystals resolved by x-ray [12].

While Spirals are generally assigned to structural functions, most catalytic disulfides are of the -/+RHHook configuration whereas the known allosteric disulfide bridges are – RHStaple. The –RHStaple, –LHHook and −/+RHHook disulfide bridges have allosteric potentials meaning that changes in such high energy bonds can influence and engender conformational changes in other parts of the said protein *(See figure 1)*. Hoggs et al. subsequently introduced –LHStaple as another form of allosteric disulfide bond, mostly encountered in crystals resolved by NMR [11]. Although it is mostly encountered in NMR crystals and rather rare in x-ray crystals, -LHStaple configuration is encountered in the TTSB Cys331-Cys418 in x-ray crystal 4R4N chains B and I *(see tables 1,2 and 3)*

**Figure 1.**
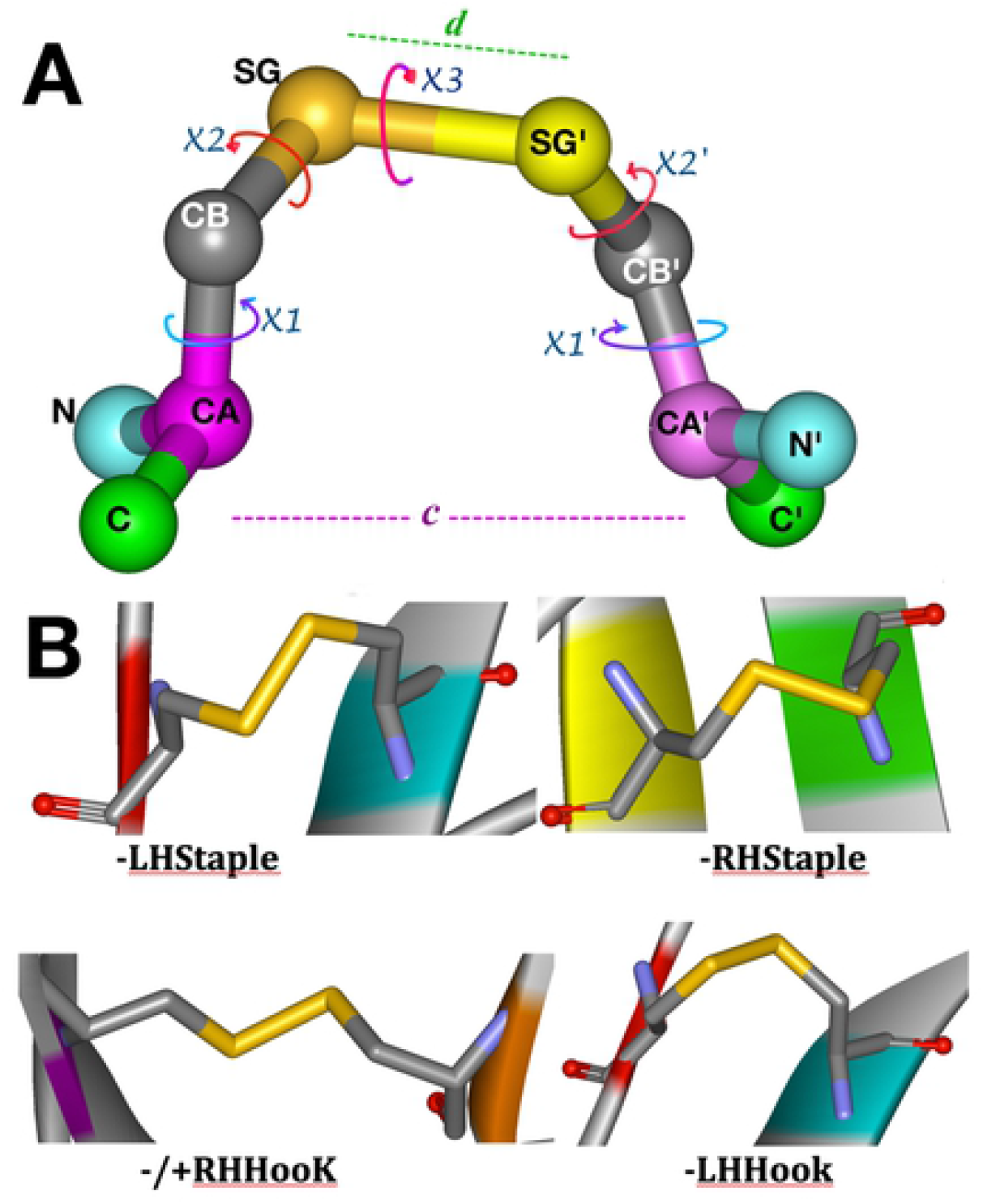
**A:** The structure of a disulfide bridge (Cys385-Cys418 in 3HI1 crystal) CA=alpha-carbon, CB= beta-carbon, SG=sulfur atom in Cys1; SG’=Sulfur atom in Cys2; *‘c’* distance between alpha-carbon atoms; *‘d’* distance between sulfur atoms. X1= chi angle between the alpha- and beta-carbon; X2= chi angle between the beta-carbon and sulfur atom; X3=chi angle between Sulfur SG and SG’ atoms of Cys1 and Cys2 respectively [11,12] **B:** Various forms of allosteric disulfide bonds. -**LHStaple** at (C418-C385) 4RFO *Chain G*; -**RHStaple** (C331-C296) 4RFO-*Chain G*; -/+**RHHooK** at C445-C378 of 2B4C *Chain G*; -**LHHook** 1RZJ (C418-C385)- *Chain G*.

**Table 1.**
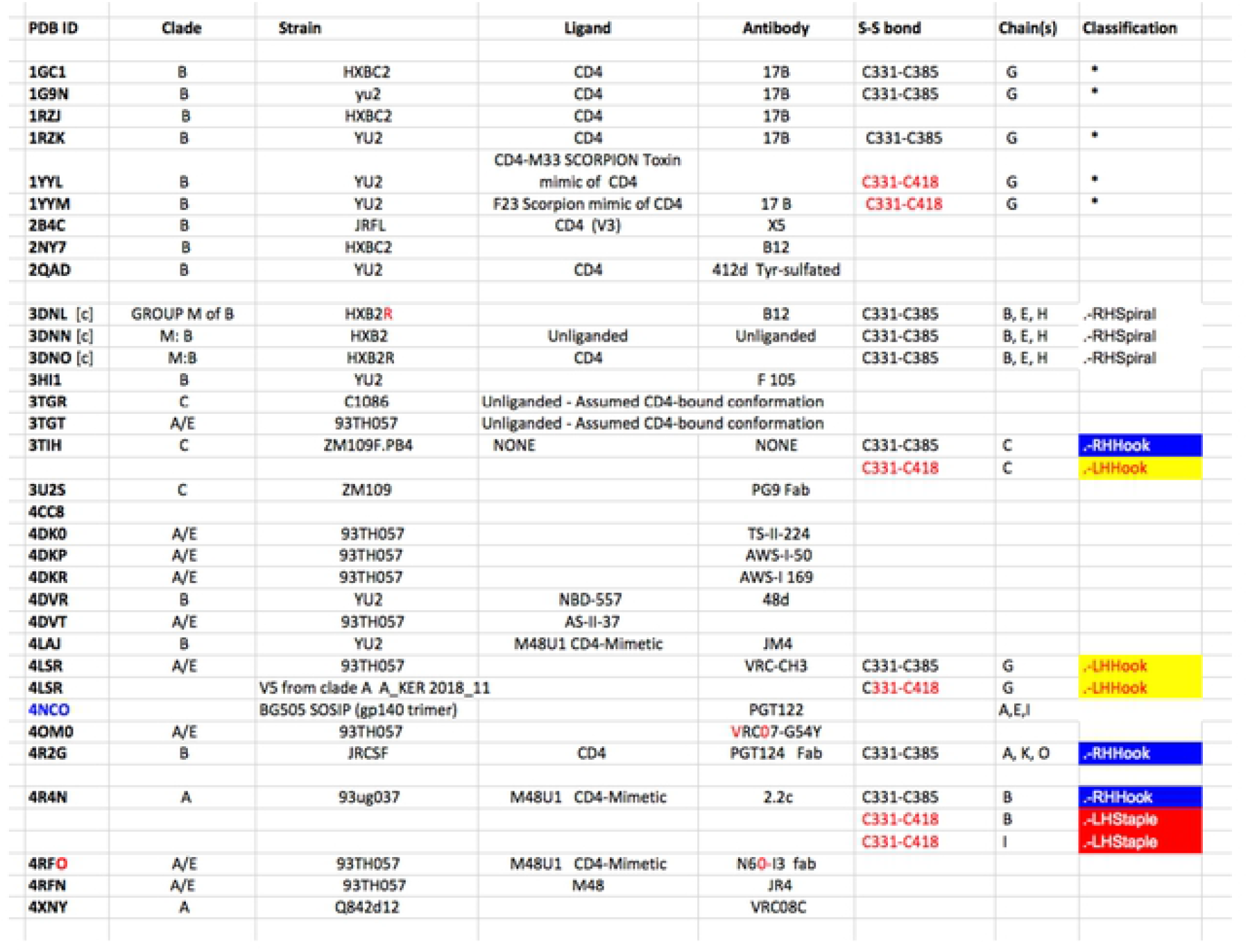
Attributes of various gp120-gp140 and gp160 crystals explored in this study. [c]: crystals resolved by Cryo-Electron Microscopy

**Table 2.**
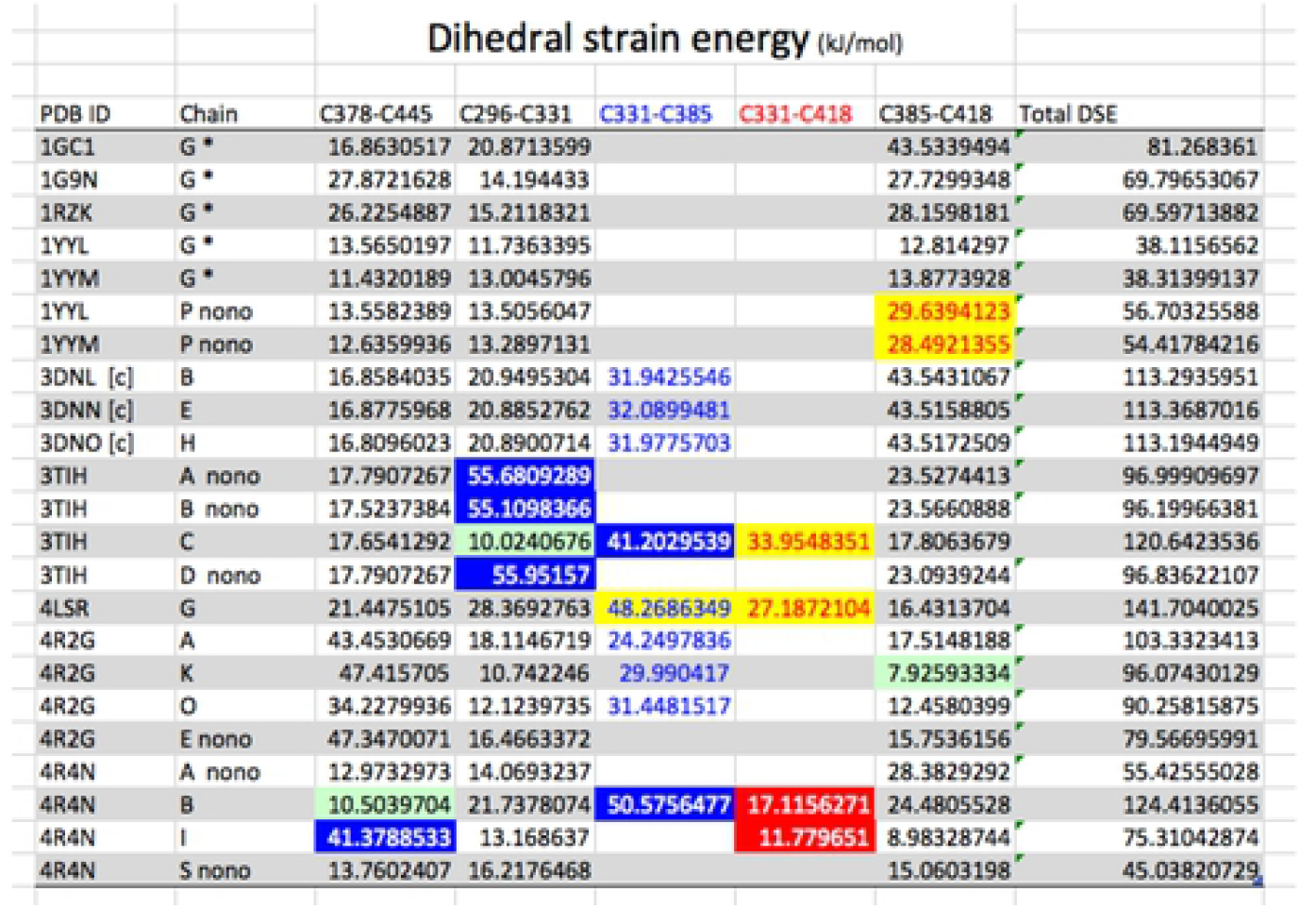
Dihedral Strain Energies *(DSE)* reported by the UNSW DBA engine. Legend: [c]= Crystal resolved by Cryo-Electron Microscopy. The other crystals are resolved by x-ray diffraction. “nono” denotes no TTSB formation in said chain. “Total DSE” pertains to the addition of the DSE’s reported by the UNSW DBA engine for the allosteric bonds involving C378, C445, C296, C331, C385 and C418 in the said chains of the said crystal. * = Chains where TTSB is observed on viewing software but not reported by DBA engine. Hence the “Total DSE” reported by the UNSW DBA engine in these *chains may miss the energy of the TTSB that are seen on the software and not reported by the DBA engine. Some key disulfide bond classifications are noted by the highlights: –RHHook, –LHHook, and –LHStaple. Most of the other allosteric bonds remained in – RHStaple configuration. Light green highlights a compensatory drop in DSE by virtue of the TTSB formation.

| Dihedral strain energy (kJ/mol) |  |  |  |  |  |  |  |
| --- | --- | --- | --- | --- | --- | --- | --- |
| PDB ID | Chain | C378-C445 | C296-C331 | C331-C385 | C331-C418 | C385-C418 | Total DSE |
| 1GC1 | G * | 16.8630517 | 20.8713599 |  |  | 43.5339494 | 81.268361 |
| 1G9N | G * | 27.8721628 | 14.194433 |  |  | 27.7299348 | 69.79653067 |
| 1RZK | G * | 26.2254887 | 15.2118321 |  |  | 28.1598181 | 69.59713882 |
| 1YYL | G * | 13.5650197 | 11.7363395 |  |  | 12.814297 | 38.1156562 |
| 1YYM | G * | 11.4320189 | 13.0045796 |  |  | 13.8773928 | 38.31399137 |
| 1YYL | P nono | 13.5582389 | 13.5056047 |  |  | 29.6394123 | 56.70325588 |
| 1YYM | P nono | 12.6359936 | 13.2897131 |  |  | 28.4921355 | 54.41784216 |
| 3DNL [c] | B | 16.8584035 | 20.9495304 | 31.9425546 |  | 43.5431067 | 113.2935951 |
| 3DNN [c] | E | 16.8775968 | 20.8852762 | 32.0899481 |  | 43.5158805 | 113.3687016 |
| 3DNO [c] | H | 16.8096023 | 20.8900714 | 31.9775703 |  | 43.5172509 | 113.1944949 |
| 3TIH | A nono | 17.7907267 | 55.6809289 |  |  | 23.5274413 | 96.99909697 |
| 3TIH | B nono | 17.5237384 | 55.1098366 |  |  | 23.5660888 | 96.19966381 |
| 3TIH | C | 17.6541292 | 10.0240676 | 41.2029539 | 33.9548351 | 17.8063679 | 120.6423536 |
| 3TIH | D nono | 17.7907267 | 55.95157 |  |  | 23.0939244 | 96.83622107 |
| 4LSR | G | 21.4475105 | 28.3692763 | 48.2686349 | 27.1872104 | 16.4313704 | 141.7040025 |
| 4R2G | A | 43.4530669 | 18.1146719 | 24.2497836 |  | 17.5148188 | 103.3323413 |
| 4R2G | K | 47.415705 | 10.742246 | 29.990417 |  | 7.92593334 | 96.07430129 |
| 4R2G | O | 34.2279936 | 12.1239735 | 31.4481517 |  | 12.4580399 | 90.25815875 |
| 4R2G | E nono | 47.3470071 | 16.4663372 |  |  | 15.7536156 | 79.56695991 |
| 4R4N | A nono | 12.9732973 | 14.0693237 |  |  | 28.3829292 | 55.42555028 |
| 4R4N | B | 10.5039704 | 21.7378074 | 50.5756477 | 17.1156271 | 24.4805528 | 124.4136055 |
| 4R4N | I | 41.3788533 | 13.168637 |  | 11.779651 | 8.98328744 | 75.31042874 |
| 4R4N | S nono | 13.7602407 | 16.2176468 |  |  | 15.0603198 | 45.03820729 |

**Table 3.**
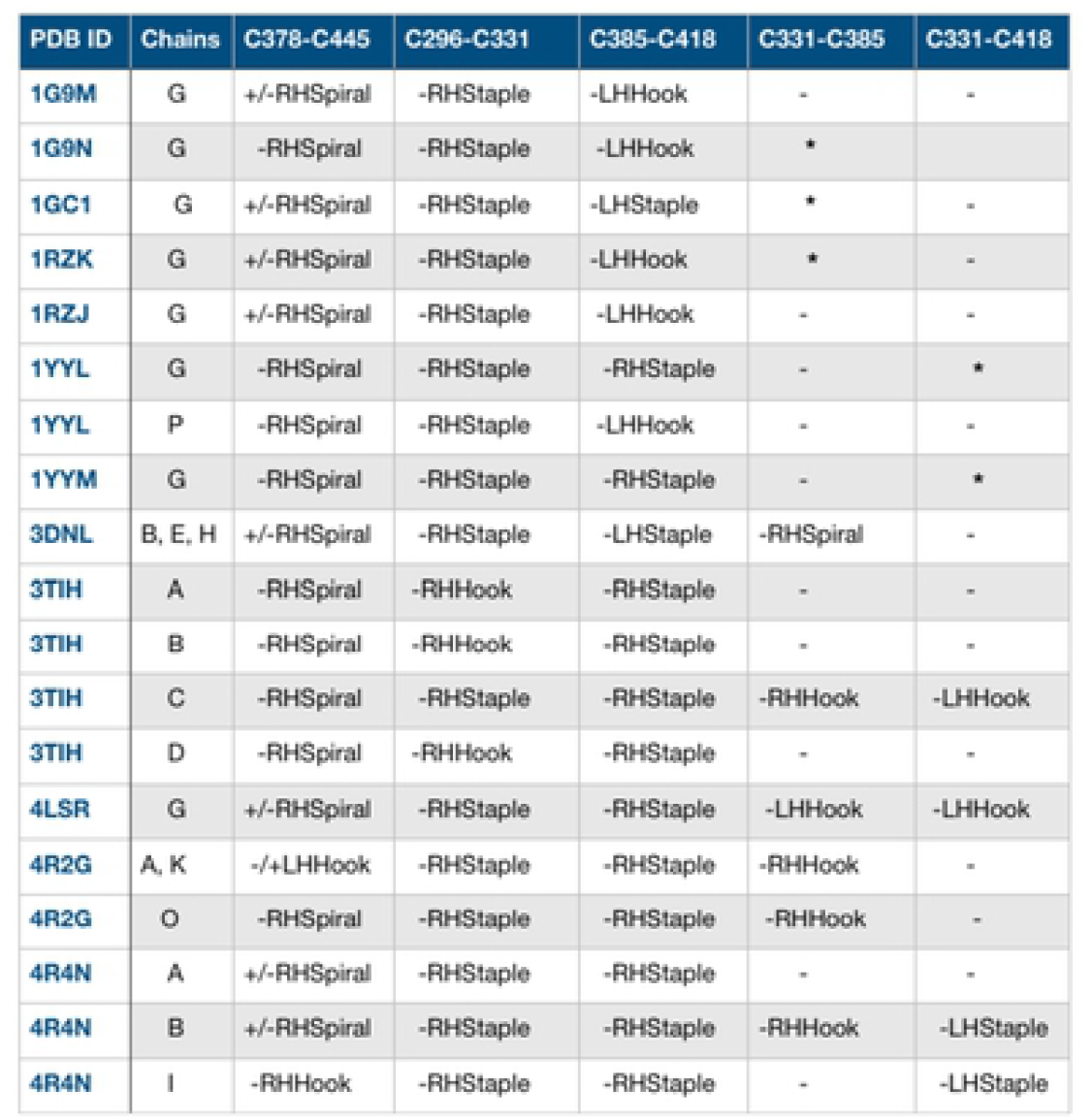
Some disulfide bond configurations. The Cys296-Cys331 disulfide bond which straddles two strands in the same beta sheet for the V3 loop mostly assumes a –RHStaple configuration except in rare conditions as seen in crystal 3TIH where it assumes a –RHHook configuration in chains A, B and D (see table 1). Cys378-Cys445 bond configuration varies between +/-RHSpiral and –RHSpiral with one instance of –RHHook in 4R4N chain I. The Cys385-Cys418 disulfide bond varies between –RHStaple, -LHHook and –LHStaple. * = Chains where TTSB is observed on viewing software but not reported by DBA engine.

The two disulfide bridges Cys296-Cys331 at the base of the V3 loop and the Cys385-Cys418 in the V4 loop are known to have significant potential for allosteric control in the gp120 molecule. On DBA analyses, they score the –RHStaple, –LHHook, −/+RHHook and –LHStaple bond types in various crystals, which places them in the classification of Cross-strand disulfides (CSDs). Cross-strand disulfides (CSDs) are unusual highly strained bonds that store torsional energy and deformation energy in the beta-sheets where they hold adjacent strands. While they are rare in usual protein structures, they are mostly encountered in proteins that are involved in cell entry (e.g botulinum toxin, HIV gp120, influenza hemagglutinin and neuraminidase [11].

The folding of gp120 is a very complex, slow but efficient process that requires multiple isomerization steps of various disulfide bridges until the correct conformation is achieved in order to allow proper cleavage of leader peptide [13]. Of note is the nature of the Cys296-Cys331 disulfide bond which straddles two strands in the same beta sheet for the V3 loop. This is a relatively rare occurrence in proteins according to Hogg [14]. The Cys296-Cys331 disulfide bond mostly assumes a –RHStaple configuration except in rare conditions as seen in crystal 3TIH where it assumes a –RHHook configuration in chains A, B and D *(see tables 1 and 3)*. Cys378-Cys445 bond configuration varies between +/- RHSpiral and –RHSpiral with one instance of –RHHook in 4R4N chain I. the Cys385-Cys418 disulfide bond varies between –RHStaple, -LHHook and –LHStaple. As noted earlier Cys418 is part of the LPCR motif in gp120 [5,6,7]

## Materials and Methods

### Gp120

The gp120 crystals in this study were downloaded from the PDB website. The gp120 molecule presents formidable difficulties for x-ray crystallography. Most crystals of gp120 are in combinations with various ligands, CD4, or antibodies *(see table 1)*. In this study we first used the gp120 structure described by Kwong et al. in 1998. The gp120 crystallized in this ternary structure was from the HXBc2 strain of HIV-1. The PDB format of the file 1GC1 for Kwong’s gp120 structure was downloaded from the PDB website. The following pdb files were subsequently studied: 1RZJ, 1RZK, 1YYL, 1YYM, 2B4C, 2NXY, 2NY0, 2NY3, 2NY4, 2NY5, 2NY7, 2QAD, 3DNL, 3DNN, 3DNO, 3G9R, 3HI1, 3NGB, 3M0D, 3P7K, 3TGQ, 3TGR, 3TGT, 3TIH, 3U2S, 3U7Y, 4DKO, 4DKP, 4DKR, 4DVR, 4DVT, 4I53, 4I54, 4JM2, 4JPW, 4JZW, 4JZZ, 4K09, 4KA2, 4LAJ, 4LSV, 4NCO, 4OLU, 4OLV, 4OLW, 4OLX, 4OLY, 4OLZ, 4OM0, 4OM1, 4NCO (SOSIP), 4R2G, 4P9H, 4RFN, 4RFO, 4RX4, 4XMP, 4XNY, 4YDI, 4YDJ, 4YE4, 4YFL,

## Software

Various studies of the gp120 molecule crystals including the mapping of key residues and measurement of bond length, bond angles, distances between residues and 3-D rendering of the gp120 molecule were achieved using the WebLab ViewerPro (version 3.7) and UCSF Chimera version 1.10.2 software packages. The WebLab ViewerPro was purchased from Molecular Simulations Inc., then Accelrys. The UCSF Chimera software was used from their website: *http://www.rbvi.ucsf.edu/chimera*. Computer-aided operations were performed on personal computers running Windows 98, XP, Windows 7 and Mac OS. The generated graphics were captured in PNG and TIFF formats and were labeled using Pixelmator.

### Disulfide Bond analyses

The gp120 crystals explored in this study were submitted through the online University of New South Wales’ UNSW Disulfide Bond Analysis engine *http://149.171.101.136/python/disulfideanalysis/search.html* (updated to https://powcs.med.unsw.edu.au/research/adult-cancer-program/services-resources/disulfide-bond-analysis-tool/disulfide-bond and https://powcs.med.unsw.edu.au/user/login?destination=node/301300173, which now requires a UNSW user account) for disulfide bond classification, chi angle measurements, dihedral strain energy (DSE) calculations, as well as distance between sulfur atoms and alpha-carbon atoms of the Cysteine residues in the respective bridges.

The 1GC1 gp120 was explored and the TTSB was observed since 1999. https://patents.google.com/patent/WO2003048186A3/en [15]. Additional crystals of gp120 were explored at random for various clades and ligands: some had TTSB formation, some did not. TTSB formation in crystals can be predicted on DBA analysis by selecting for “331” in the Cys 1 position on the spreadsheet because Cys331 is usually in Cys 2 position with Cys296 in the Cys 1 position whereas in position 1 it relates to either Cys385 and/or Cys418 connections as is discussed below. All the TTSB’s reported on the DBA analysis -including the (CYS331:SG –CYS385:SG) and the CYS385:SG - CYS418:SG and the combined TTSB as observed in 3TIH – were also observed in the viewing software. However some TTSB’s like the ones in 1GC1 and 1RZK crystals seen on the viewing software were not detected or reported by the DBA analyses.

## Results

### Viewing Software analyses

*Exploring the 1GC1 crystal chain G*, looking at the outer domain *(shell view)* of the gp120 molecule *(see figure 2 A)*, we observe a beta-pleated sheet with three strands spanning the central area. Each of these strands contains a cysteine residue actually juxtaposed in the same plane in the 3-dimensional configuration. In figure 2, these Cysteine residues are from left to right: #1=Cys445, #2 = Cys296 and #3 = Cys331. In another plane behind these 3 juxtaposed cysteine residues lies another Beta sheet that runs almost perpendicular to the first one. There are 2 other cysteine residues in this second sheet, namely #4 = Cys385 and #5 = Cys378. Then on the side, between Cys331 and Cys385, runs another string of amino acids, *(leading to the segment assigned to the flip/switch phenomenon 417 – 429)* which contains #6 = Cys418.

**Figure 2:**
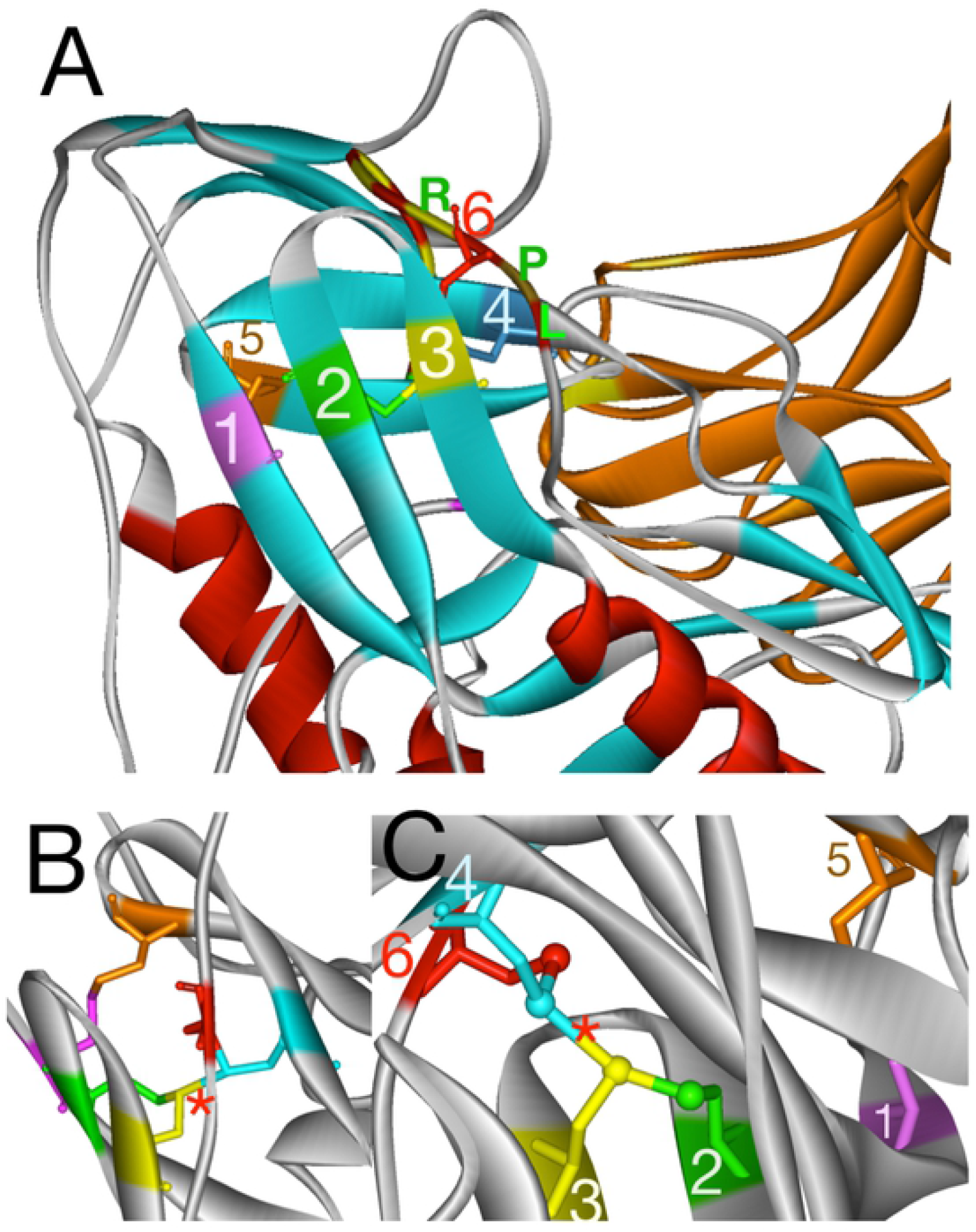
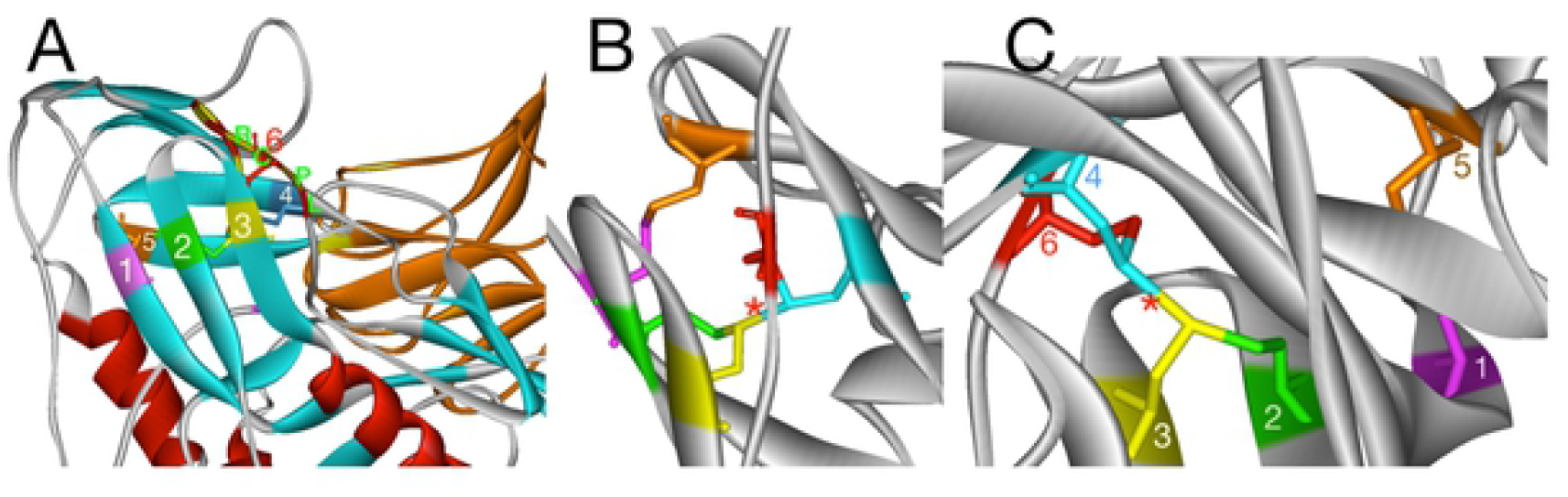
TTSB in 1GC1. **A)** Shell (outer) view of the gp120 molecule. Locations of Cystine residues in the different planes. 1=Cys445; 2=Cys296; 3=Cys331; 4=Cys385; 5=Cys378; 6=Cys418. 1 Magenta=Cys445; 2 Green=Cys296; 3 Yellow=Cys331; 4 Cyan=Cys385; 5 Brown=Cys378; 6 Red=Cys418.The LPCR sequence denotes the LPCR motif beginning of the segment *(L416 through W427)* involved in the bistable flip/switch phenomenon. **B)** 90 degree rotation showing the TTSB from the Right side view. The red asterisk *denotes the TTSB. The disulfide bridge (C445-C378) and the tetrasulfide bridge (*TTSB) keep the two beta sheets together. **C)** Ventral view of the TTSB

#### Exploring the 1GC1 crystal

Analysis of the gp120 structure in 1GC1 crystal using the WebLab ViewerPro and UCSF Chimera software reveals the following sulfide bridges *(see figures 2)* between the above mentioned cysteine residues:

Cys378 makes a disulfide bridge with Cys445 = Bridge #1
Cys296 makes a disulfide bridge with Cys331 = Bridge #2 for the V3 loop
Cys385 makes a disulfide bridge with Cys418 = Bridge #3 for the V4 loop

From bridge #2 and Bridge #3, the sulfur atom from Cys331 shares another bond with the sulfur atom of Cys385 forming a new CONNECTION (CYS331:SG – CYS385:SG) that links the CYS296:SG - CYS331:SG bridge with the CYS385:SG - CYS418:SG bridge. This new connection forms a new bridge between two disulfide bridges. It is the CYS331:SG – CYS385:SG bond.

Given the fact that the CYS331:SG – CYS385:SG bond is a bridge formed by 4 sulfur molecule arrangement, this novel type of bond is henceforth referred to as a tetrasulfide bridge (or TTSB) in the rest of the article. This is the first time such a bridge is being described in protein. While TTSB’s may well exist undiscovered in other proteins, this CYS331:SG – CYS385:SG bond represents a unique landmark in the architecture of the gp120 of HIV-1

##### Exploring other gp120 crystals

Analysis of other gp120 crystals from various HIV strains in various combinations of ligands and antibodies reveal mixed results *(see figures 2–6)*. The crystals were viewed using various rendering software Weblab Pro Viewer and UCSF Chimera and both software showed the TTSB in the respective crystals. The TTSB formation was noted in various chains of the following crystals: 1GC1, 1G9N, 1RZK, 1YYL, 1YYM, 3TIH, 3DNL, 3DNN, 3DNO, 4LSR, 4R2G, 4R4N *(see figures 4 and 5)*.

TTSB formation was observed in crystals 1GC1 chain G, 1G9N chain G, 1YYL chain G but not chain P, 1YYM chain G but not chain P, 4R2G clade B in chains A, K and O but not in chain E; 1RZK chain G, all chains B, E and H of trimers 3DNO, 3DNL and 3DNN and in chain C but not in chains A, B nor D of the unliganded 3TIH crystal of a clade C strain. TTSB formation was also noted in crystals 4LSR chain G, 4R4N chains B and I. TTSB formation was absent in the other PDB crystals analyzed by the above software in this study. In the 1YYL and 1YYM chains G crystal, as well as 4R4N chain I, the TTSB was demonstrated in both software to involve the SG of the Cys418 as opposed to the SG-Cys385 which is the most common configuration *(see figure 5)*. Chain C of 3TIH, chain G of 4LSR as well as chain B of 4R4N crystals of gp120 show a triangular form of TTSB intermediate with <u>both</u> the TTSB (Cys331-Cys385) and the Cys331-Cys418 *(see figure 6)*. Coil helix form was not noted in the 418-427 segment in any of the crystals explored in this study irrespective of TTSB formation.

### UNSW DBA Analyses

DBA analysis reported the presence of TTSB formation in some but not all of the crystals where the viewing software Weblab Pro and UCSF Chimera had shown them. In crystals resolved by X-ray diffraction, the DBA analysis detected TTSB (Cys331-Cys385) in 4RG2 chains A, K,O, as well the TTSB (Cys331-Cys385) with the Cys331-Cys418 intermediate bond in chain C of 3TIH, chain B of 4R4N and chain G of 4LSR crystals *(see figure 6)*. DBA analysis did report Cys331-Cys-418 TTSB formation in 4LSR chain G and 4R4N chain I; but the DBA analysis did not detect the Cys331-Cys-418 TTSB formation observed in the 1YYL and 1YYM crystals in the viewing software. In the crystals resolved by Cryo-Electron Microscopy, the DBA analysis reports the TTSB in this study in chains B, E and H of 3DNN, 3DNL, 3DNO all of which pertaining to the trimeric forms of gp120 where 3DNN is unliganded, 3DNL is bound to b12 antibody and 3DNO bound to CD4 *(see table 1)*.

The presence of the TTSB could be affected by the target domains of the various antibodies used in the crystallization processes. In order to assess the degree of deformity or distortion of the gp120 in various crystals, we monitored 2 distances in 3 conserved residues: D1= distance between CG2 of Thr257 – and CZ3 of Trp427 in Angstroms and D2= the distance between CG2 of Thr257 and CB of Glu370 *(see figure 8)*. There did not seem to be any correlation between TTSB formation and D1 or D2 measurements (data not shown) Range for Crystals with TTSB: 9.72A- to 10.43A for D1 and between 4.73A – 5.04A for D2. Whereas in crystals without visible TTSB formation on utilized software D1 ranged from 10.09A to 10.23A and D2 between 4.46 and 5.02A. Noteworthy is the fact that chain G of 3HI1 crystal *(not 3TIH but 3HI1)* -which showed no TTSB formation and which represents gp120 bound to CD4-binding site antibody F105 - had D1 of 27.33A *(which reflects major deformation of the gp120)* and a D2 of 4.48A. As illustrated in table 2, there are major changes in dihedral strain energies (DSE) in the allosteric bonds involved in TTSB formation. More importantly the data reveal a shift of DSE’s between the allosteric bonds within the respective chains with TTSB formation.

### Partial charges

Partial charges in sulfur atoms in disulfide bonds is generally −1. However in the triangular shaped TTSB intermediate formation [3TIH chain C, 4LSR chain G and 4R4N chain B] the sulfur atom in Cys331 assumes a sulfonium characteristic with a +positive partial charge owing to the high nucleophilicity of sulfur *(see table 4)*.

**Table 4.**
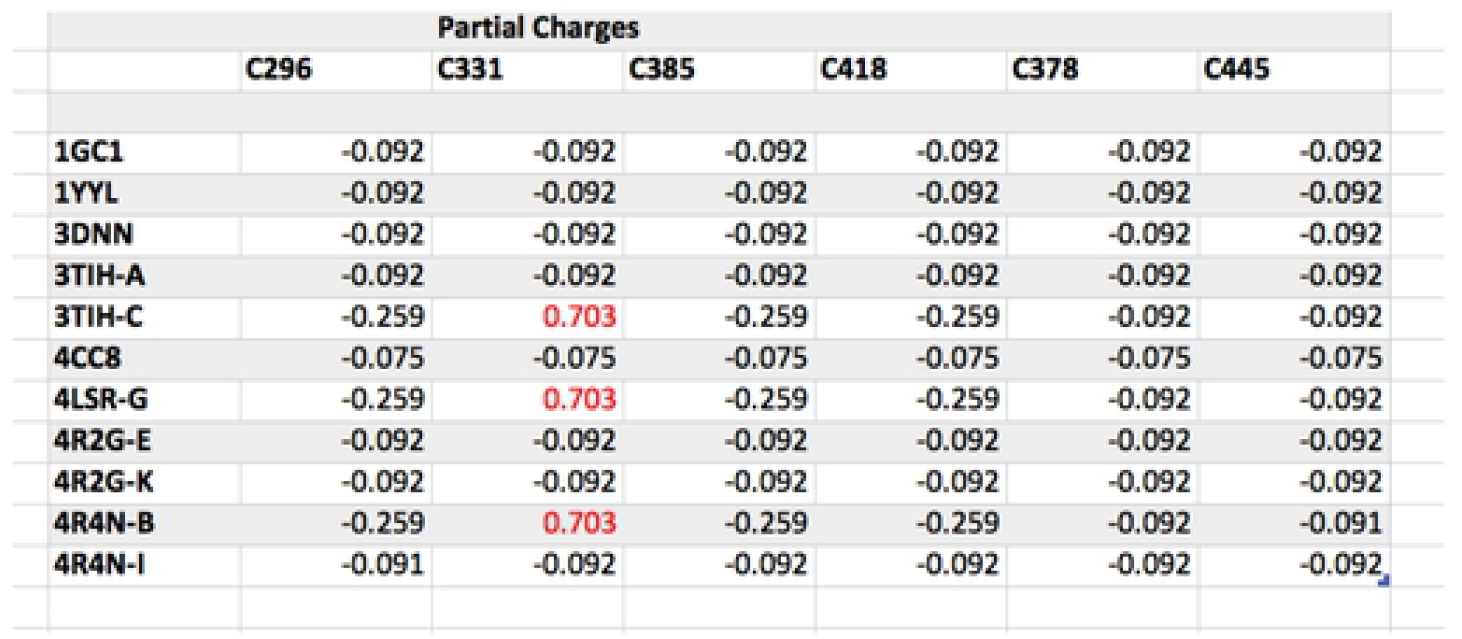
Partial charges of sulfur atoms in TTSB formations. – as monitored in WebLab Pro software.

Note in triangular shaped TTSB formation [3TIH chain C, 4LSR chain G and 4R4N chain B] the Sulfur atom in Cys331 assumes a +positive partial charge.

Noteworthy are the striking differences in characteristics of some allosteric bonds in the various configurations:

### C331-C385 and C331-C418 TTSB’s

The TTSB formations observed in this study mostly occur as Cys331-Cys385; but in Chains G of 1YYL, 1YYM crystals, chain G of 4LSR and chain I of 4R4N, it exists as Cys331-Cys418. Whereas in Crystal 3TIH chain C, 4R4N chain G, 4LSR chain G, it exists as both Cys331-Cys385 and Cys331-Cys418 *(See figures 4 and 5)*

### Sulfur atom distances

TTSB bond length measurements (i.e. Distance between sulfur atoms of the Cys residues in Å(Angstroms) by viewing software and/or by DBA reports fall between 2.734 and 2.998 Å, except for the Cys331-Cys385 and Cys331-Cys418 in crystal 3TIH that measured the shortest with 2.092 and 2.104 Angstroms respectively, although the ones in 4R2G the Cys331-Cys385 TTSB measured 2.53, 2.94 and 2.57 A in chains A, K and O respectively.

### Carbon atom distances

Most disulfide bridges reported in our DBA analysis have a Distance between α-carbon atoms of the Cys residues, (Å) that fall between 3.6 and 4.3Angstroms whereas the TTSB that distance falls between 6.865 and 7.19 A. In the Cys331-Cys418 bridge in crystal 3TIH it measured 6.355A.

### DSE (dihedral strain energy) changes in the C296-C331 bond

*(See table 2)*. Based on thiol transfer dynamics, the higher the strain energy the more likely for the bond to be cleaved [14, 16, 17]. In the 3TIH crystals in chains A, B and D which do not show any TTSB formation, the dihedral strain energy (DSE) for the Cys296-Cys331 bridge score the highest in our study: 55.681 kJ/mol, 55.1098 and 55.951 kJ/mol respectively with a - RHHOOK configuration as opposed to its usual -RHStaple characterization.. Whereas in 3TIH chain C which has both Cys331-Cys385 with Cys331-Cys418 TTSB in the triangular configurations *(see figure 6)*, the dihedral strain energy for the Cys296-Cys331 bridge score the lowest in our study: 10.024 kJ/mol where it still maintains a –RHStaple characteristic *(see tables 2 and 3)*.

### DSE changes in the C378-C445 bond

A similar drop in DSE to 10.504 Kj/mol is also observed in **C378-C445** bond in crystal 4R4N chain B which also has a triangular shaped TTSB with high DSE values (of 50.576 kj/mol in the C331-C385) in the TTSB formation. Whereas the energy is increased to 41.379 kj/mol with –RHHook configuration in the C378-C445 bond in chain I of crystal 4R4N which has a C331-C418 TTSB formation.

### DSE for TTSBs

In 3DNN, 3DNL and 3DNO crystals, DSE for TTSB (C331-C385) range from 31.942 and 32.089 in a –RHSpiral confiruration *(see tables 1 and 2)*. In the 4R2G crystal the (C331-C385) TTSB records DSE’s of 24.249, 29.990 and 31.448 kJ/mol with a –RHHook character for chains A, K and O respectively. In the 3TIH crystal chain C which has the triangular TTSB formation, the Cys-331-Cys385 TTSB absorbs a DSE of 41.203 kJ/mol with a –RHHook configuration while the Cys331-Cys418 bond scores 33.955 kJ/mol with a -LHHook. As noted above, while the TTSB’s in the 3TIH chain C have high DSE values, the Cys-296-Cys331 and the Cys385-Cys418 allosteric bonds drop their DSE scores to 10 kJ/mol and 17.8 kJ/mol respectively. That suggests a safe transfer of energy between the bonds thanks to the TTSB formation.

#### 4R4N

In crystal 4R4N chain I, the TTSB (C331-C418) had a DSE of 11.78 kj/mol while the C378-C445 bond had a DSE of 41.38 kj/mol in the same chain I; whereas the DSE in C378-C445 bond of in 4R4N chain B which has a triangular TTSB formation drops to 10.50 kj/mol with +/-RHSpiral configuration. In the same chain B of 4R4N there is a compensatory DSE absorption of 50.58 kJ/mol in the (C331-C385) TTSB bond and 17.12 kj/mol in the (C331-C418) TTSB bond.

As noted above, crystals 1YYM chain G and 1YYL chain G had TTSB with (Cys 331-Cys 418) TTSB bond observed in viewing software but not reported in the DBA analyses. The same for crystals 1GC1, 1G9N and 1RZK had TTSB with (Cys331-C385) bond observed in viewing software but not reported in the DBA analyses.

### Energy transfer between the allosteric bonds

The changes in energy observed in the allosteric bonds in table 2 illustrate a flux of energy transfer between the allosteric bonds C378-C445, C296-C331 and C385-C418 during TTSB formation. These changes are more extreme in C378-C445 and C296-C331 bonds. The extreme changes in dihedral strain energy (DSE) suggest both protection and exposure of the C378-C445 and C296-C331 bonds to thiolate attacks.

Disulfide bond cleavage is an important step in the process of gp120 binding to CD4. In fact as illustrated by Gallina et al, PDI inhibitors prevent HIV entry in CD4+ cells [16]. Azimi et al demonstrated that the Cys-296-Cys331 bond is cleaved by thioredoxin [17, 18]. This is understandable when this bond achieves high DSE. However during the formation of the TTSB intermediate illustrated in unliganded crystal 3TIH chain C, the DSE for the Cys296 bridge goes to its lowest (10kj/mol). Based on thiol transfer dynamics [16], that would likely protect the Cys296 against thiolate attacks in 3TIH chain C whereas in 3TIH chains A, B and D which have no TTSB formation the DSE for the C296-C331 bond goes up to over 55kj/mole, which could expose the bond to thiolate attack. **The sequence of these events cannot be explained with the available data in our study.** 4R4N chains B & I illustrate a dynamic transfer of energy between the allosteric bonds in this study thanks to the TTSB formation.

The TTSB formation may absorb energy to protect the C296-C331 bond for some time (eg before CD4-binding, as seen in the unbounded gp120 crystal of 3TIH) in order to expose it at a later time to thiolate attack after CD4-binding. The binding of CD4 favors the cleavage of the C296-C331 disulfide bond (Azimi) [17]. Whether the TTSB formation protects or exposes the C296 to thiolate attack may well depend on whether it is bound to CD4 or not. **The DSE transfers in the TTSB may program the proper timing for cleavage of the Cys296-Cys331 bond by thioredoxin to occur after CD4 binding.**

The data reveals the existence of a tetrasulfide bridge (TTSB) in the architecture of the gp120 molecule of HIV-1. Analyses of multiple crystals of gp120 suggest the existence of the TTSB more as an intermediate as opposed to a constant landmark, which implies more complex functions than just structural attributes. As illustrated in table 2 there is a transfer of DSE’s shifted between the allosteric bonds in TTSB formation.

## Discussion

Disulfide bridges are well known for their ability to add integrity to protein structures. Cross-strand disulfides (CSDs) may store kinetic energy in their highly strained bonds for dynamic functions like cell entry [11, 12, 14]. Tetrasulfide bridges on the other hand were never described before in any protein structure. The gp120 molecule has many properties that are unique to HIV. Despite the high mutation rate, the virus manages to uphold enough integrity in its 3-dimensional structure to maintain key functions such as CD4-binding, CCKR5-binding and infectivity. Using a tetrasulfide bridge in the architecture of the gp120 is a rather practical way for the virus to maintain function and integrity and also limit space, mass and volume at the same time.

As demonstrated in figure 2 and figure 3, two beta sheets in different planes are kept approximated by one disulfide bridge (Cys378-Cys445) and one tetrasulfide bridge (CYS331:SG – CYS385:SG) connecting the planes of the V3 and V4 loops. This tetrasulfide bridge did not happen by accident, nor could it be the side effect or artifacts of experimental manipulations and/or chemical exposure during the crystallization processes. TTSB formation is only observed between Cys296-Cys331 and Cys385-Cys418 bridges, namely at the bases of the V3 loop and the V4 loop. Although the Cys378-Cys445 undergoes major change in DSE potentials during TTSB formation in the gp120 molecule, no TTSB formation is observed in that bond. TTSB structure was observed in crystals of various strains from different clades of HIV with various ligands. Moreover it was also rendered visible and labeled appropriately by different viewing software and it is also reported, measured and characterized in various crystals through DBA analysis. Based on the results in our study, the TTSB appears to be an important intermediate in the dynamics of gp120 especially where CD4 binding and viral entry are concerned.

**Figure 3.**
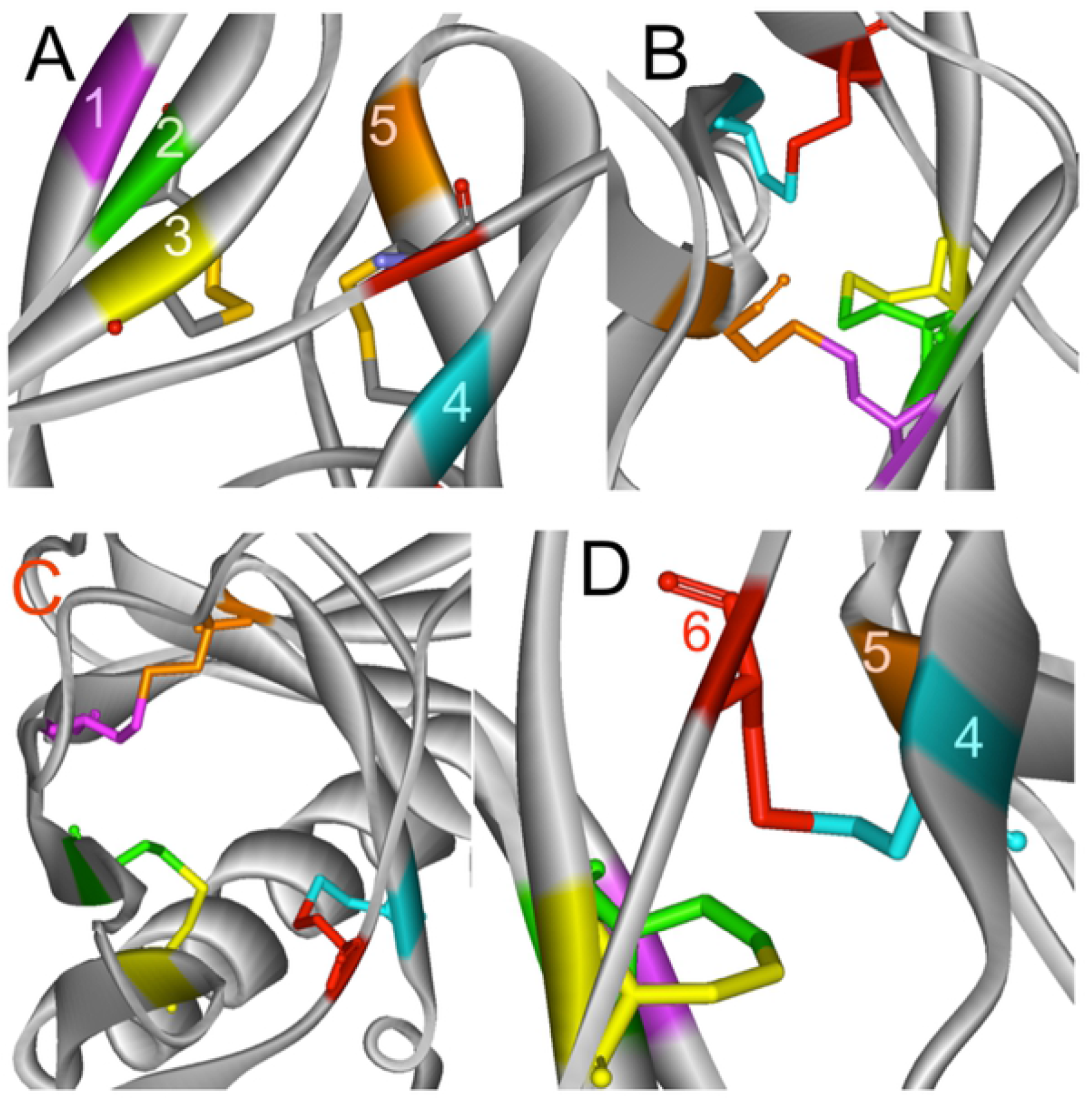
Some gp120 crystals without TTSB formation. A) 4RG2 chain E with element colors in disulfide bridges. B) 2B4C chain G Left 90 degree rotation view in coded colors. C) 3HI1 chain G top view. D) 3TIH chain A Right sided view. *Legend: 1 Magenta=Cys445; 2 Green=Cys296; 3 Yellow=Cys331; 4 Cyan=Cys385; 5 Brown=Cys378; 6 Red=Cys418*.

**Figure 4.**
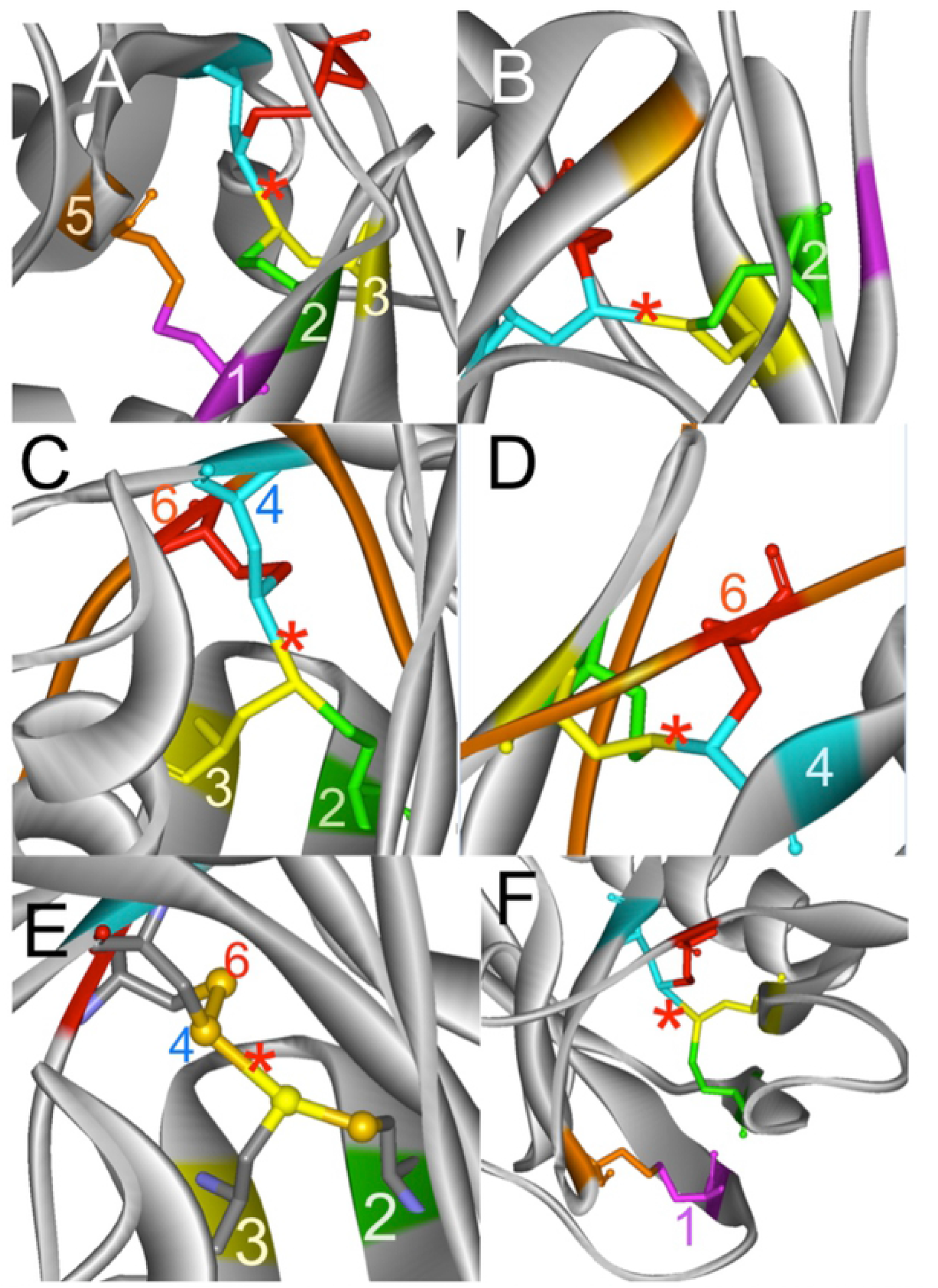
Various TTSB formation denoted by the red asterisk * with Cys331-Cys385. A) 1RZK 90 degree rotation in Left sided view, B) 1G9N, C) 3DNO, D) 3DNL. E) Ventral view of 1GC1 with element color and sulfur atoms in ball & stick representation. F) 4RG2 chain K top view. *Legend: 1 Magenta=Cys445; 2 Green=Cys296; 3 Yellow=Cys331; 4 Cyan=Cys385; 5 Brown=Cys378; 6 Red=Cys418*.

**Figure 5.**
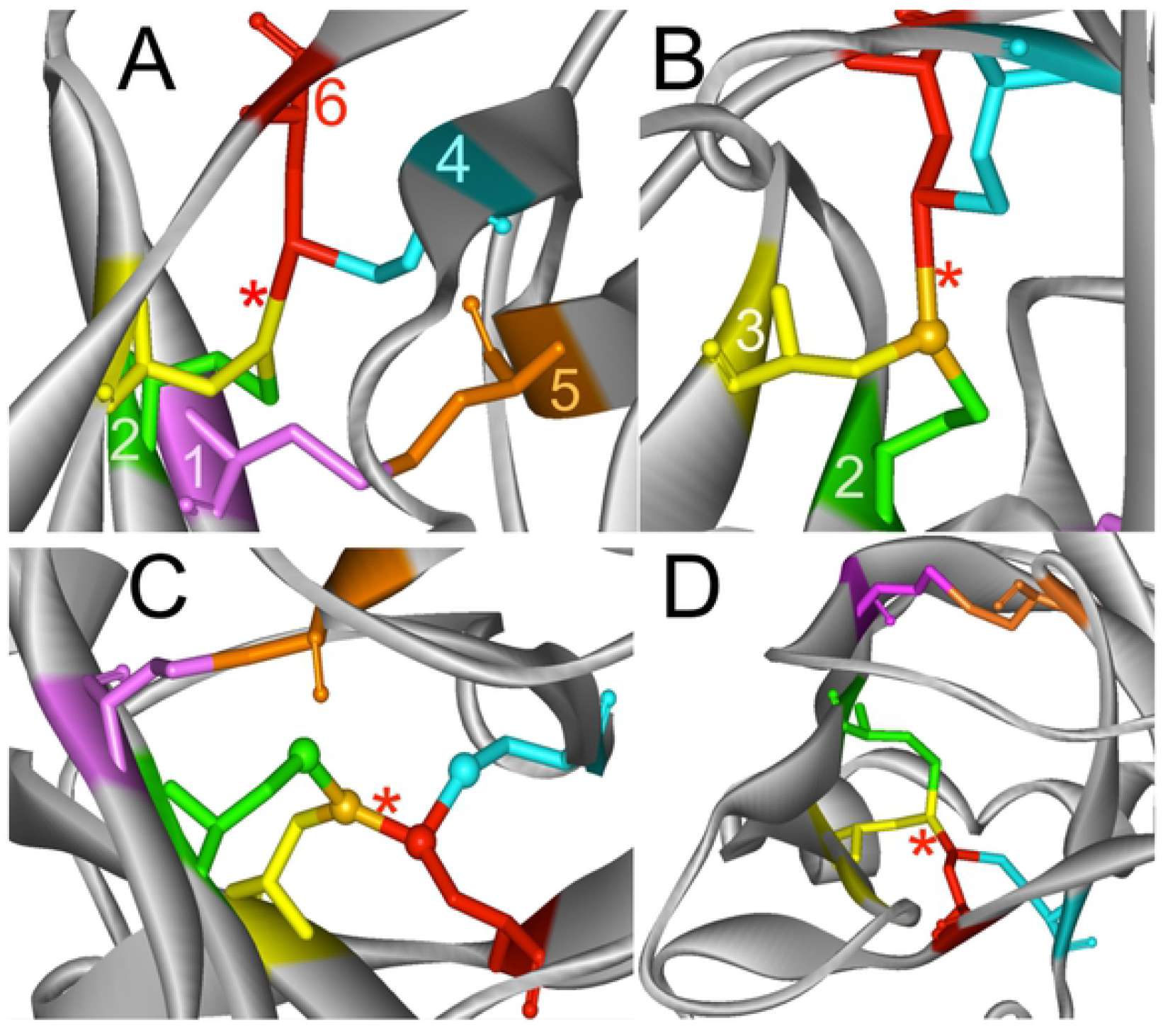
TTSB formations with red Cys418 and yellow Cys331 (Cys331-Cys418 bond). The red asterisk denotes the TTSB. A= 1YYM chain G, B= 1YYL chain G, C= 4R4N Chain I, D=4LSR chain G top view. *Legend: 1 Magenta=Cys445; 2 Green=Cys296; 3 Yellow=Cys331; 4 Cyan=Cys385; 5 Brown=Cys378; 6 Red=Cys418*.

In 1YYL, Cys385-Cys418 is –RHStaple in chain G and –LHHook in chain P. This bond assumes an –LHStaple configuration in 1GC1. The variations observed in torsional energies in these allosteric bonds are beyond the scope of this study. However the transfer of energies between the allosteric bonds explored in this study attest to a dynamic state of these bonds within the molecule. In most crystals we explored in this study the TTSB occurs as Cys331:SG – Cys385:SG single bond whereas in 1YYL and 1YYM chain G, 4R4N chain I and 4LSR chain G, it occurs as Cys331:SG – Cys418:SG bond. So TTSB can be formed combining Cys331 either with Cys385 or Cys418. Cys296 is only found bounded to Cys331. Noteworthy is the fact that in the same crystallization process for 1YYL and 1YYM, chain G has TTSB formation whereas chain P does not; as well as in crystal 3TIH where TTSB formation is found in chain C and not in chains A, B nor D. The fact that some crystals show the TTSB formation in some chains and not in other chains suggests that the TTSB may exist in an intermediate state in an allosteric position for both structural and possibly chemical/mechanical dynamics in the gp120 molecule. The striking observation in these alternating configurations with and without TTSB formation in these crystals remains the redistribution of DSE energies in these various allosteric disulfide bonds involved *(see tables 2 and 3)*.

### Triangular TTSB formation

The intermediate nature of the TTSB dynamics is best illustrated in chain C of crystal 3TIH where Cys331-Cys385 is actually reported as a – RHHook bond (on UNSW DBA analysis) while the crystal shows and a –LHHooK bond between Cys331 and Cys418 although the Cys296-Cys331 remains stable. This results in a triangular shape of TTSB between the SG sulfur atoms of Cys331, Cys385 and Cys418 *(See figure 6)*. This triangular shape of the TTSB represents a dynamic intermediate between the common TTSB between Cys331-Cys385 as seen in 1GC1 and the formation with Cys331-Cys418 configuration as observed in chain G of 1YYL and 1YYM crystals. In figure 6, also note that the SG sulfur atom of Cys331 *(yellow)* still maintains covalent bonds with its CB carbon and the SG sulfur of Cys296 *(green)*. This puts the SG sulfur atom of Cys331 in the center of a tetrahedral molecular geometry bonding with: 1) the CB carbon atom of Cys331; 2) the SG sulfur atom Cys296; 3) the SG sulfur atom Cys385; 4) the SG sulfur atom Cys418. *(See Figure 6)*. In this configuration the sulfur atom in Cys331 acquires a +positive partial charge *(see table 4)*, which favors thiol-disulfide exchange reaction within the TTSB intermediate.

**Figure 6.**
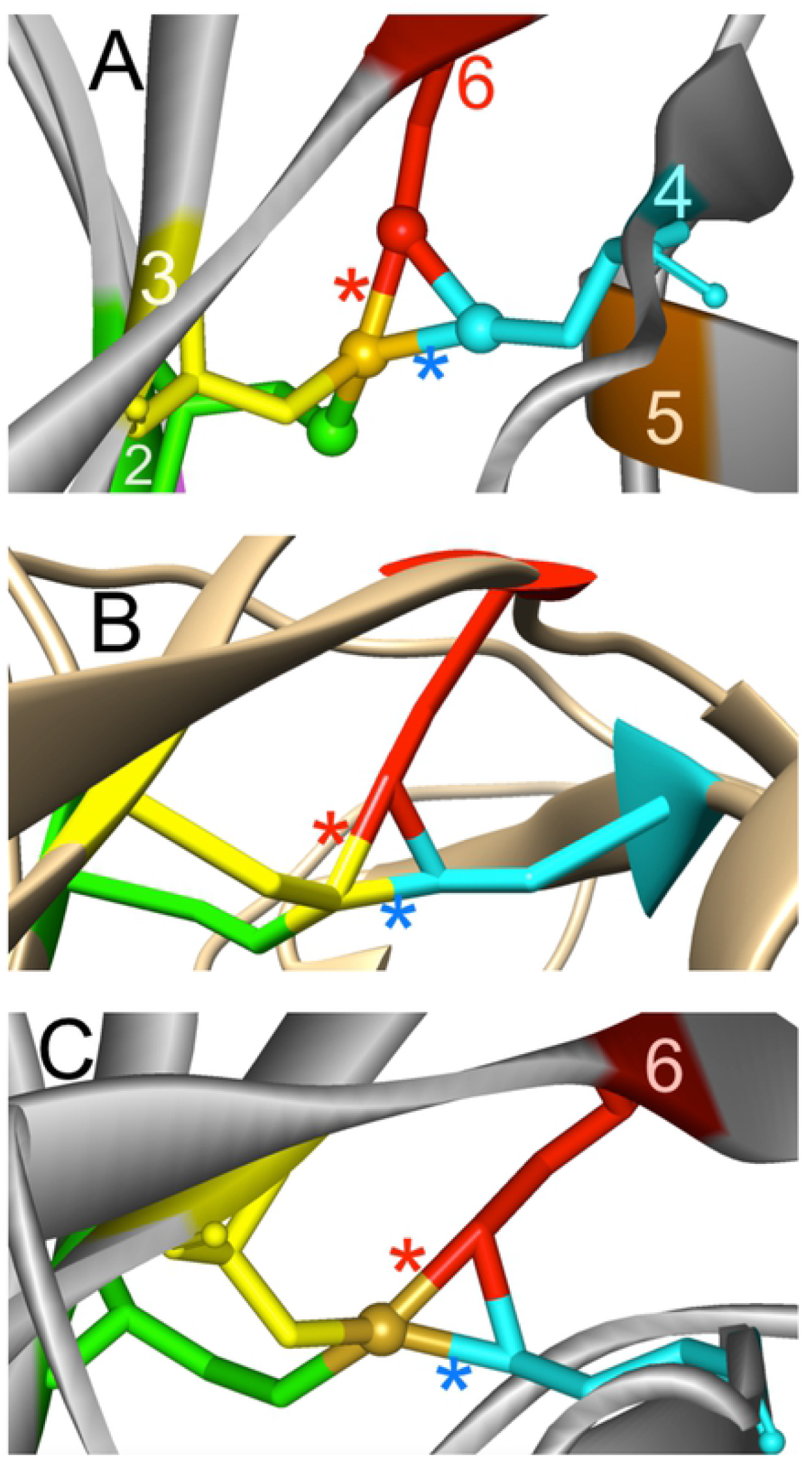
Triangular shaped TTSB intermediate formation, **A)**4R4N chain B, **B)**3TIH chain C *(as viewed in UCSF Chimera)* and **C)** 4LSR chain G. The red asterisk * denotes the Cys331-Cys418 bond and the blue asterisk * denotes the Cys 331-Cys385 bond. This triangular shape of TTSB formation seems to represent an intermediate step between the Cys331-Cys385 TTSB seen in figure 4 and the Cys331-Cys418 TTSB formations seen in figure 5. *Legend: 1 Magenta=Cys445; 2 Green=Cys296; 3 Yellow=Cys331; 4 Cyan=Cys385; 5 Brown=Cys378; 6 Red=Cys418*.

**Figure 7.**
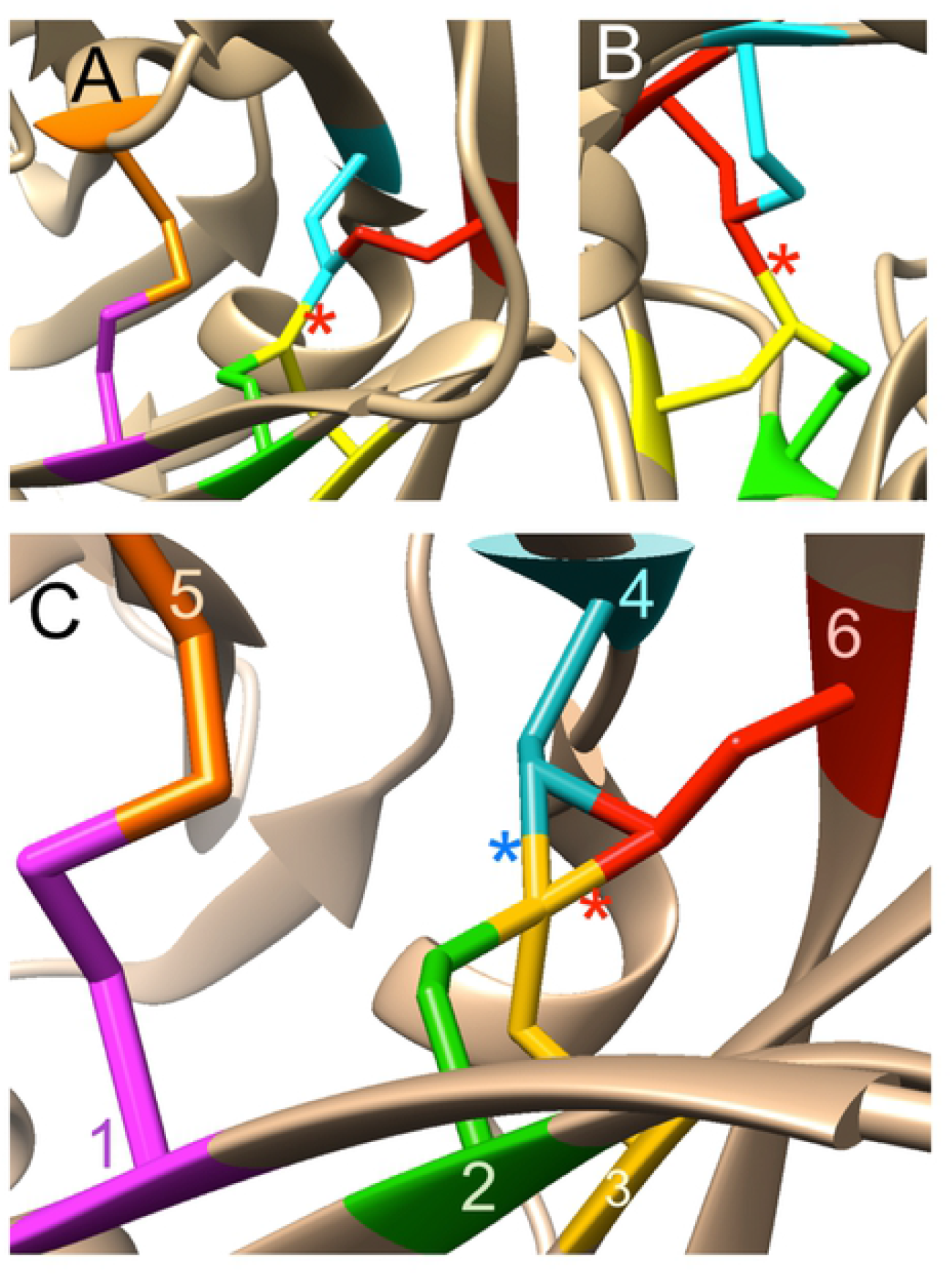
TTSB’s viewed in UCSF Chimera software. A) 1GC1 with Cys331-Cys385 TTSB, B) 1YYL with Cys331-Cys418 TTSB. C) 4LSR with triangular shape TTSB intermediate., *Legend: 1 Magenta=Cys445; 2 Green=Cys296; 3 Yellow=Cys331; 4 Cyan=Cys385; 5 Brown=Cys378; 6 Red=Cys418*.

**Figure 8.**
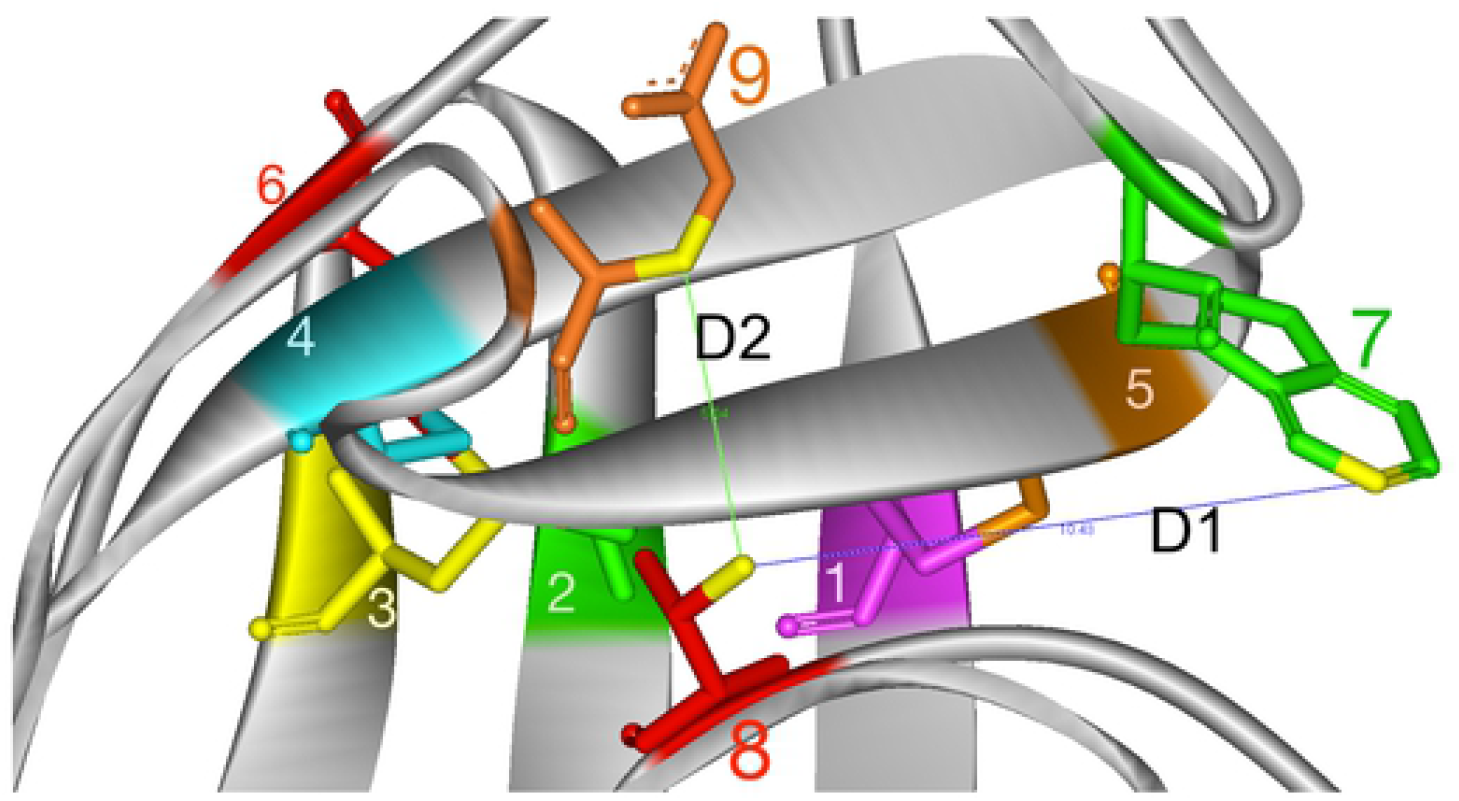
D1 and D2 measurements in 1YYL crystal. *Legend: 1 Magenta=Cys445; 2 Green=Cys296; 3 Yellow=Cys331; 4 Cyan=Cys385; 5 Brown=Cys378; 6 Red=Cys418. 7 green Trp 427; 8 red=Thr257; 9 brown=Glu370*.

The TTSB formation is similar in some way, yet very different than the expected disulfide shuffling which naturally occurs via intra-protein thiol–disulfide exchange reactions. The similarities begin with the formation of extra bonds and the breakage sulfide bonds; but the differences lie in the fact that the bonds that are broken in this case are only transient intermediates of the TTSB formation [namely Cys-331-Cys385 and Cys331-Cys418] whereas the original allosteric bonds [Cys331-Cys296, Cys378-Cys445 and Cys385-Cys418] remain unbroken in the crystals we explored although major changes do occur in their DSE values *(see table 2)*.

There is an “unknown second disulfide bond” expected to break in the thiol exchange by thioredoxin on gp120, which Schmidt and Hogg predicted to be the C385-C418 bond [12]. This “unknown second disulfide bond” may very well be dictated by TTSB intermediate formation.

#### Timing program

Reviewing tables 1 and 2, we note some interesting differences in two crystals with triangular TTSB, namely the 3TIH crystal which is of unliganded gp120 and 4R4N where gp120 is bound to a CD4 mimetic. In 3TIH chain C, the TTSB formation transfers DSE energy into the Cys331-Cys385 bond while it drops the energy in the cys296-cys331 from 55 kj/mol observed in chains A, B and D without TTSB to 10kj/mol in chain C with the triangular TTSB formation. Whereas in chain B of the 4R4N crystal where gp120 is bound to a CD4-mimetic, it transfers the energy between the Cys378-Cys445 and the transient Cys331-Cys385 TTSB. The triangular TTSB formation leads to a major reduction in DSE at the Cys296-Cys331 bond in the unliganded 3TIH chain C whereas triangular TTSB formation leads to a relative rise in DSE at the Cys296-Cys331 bond in 4R4N chain B bound to CD4-mimetic. These events represent in vitro snap shots of the phenomenon of TTSB intermediate formation. Based on thiol transfer dynamics [13, 17], the drop in DSE of the bond would protect it against thiolate attack. TTSB formation engenders DSE redistribution that seems to protect the Cys-296-Cys331 bond before CD4 binding *(as seen in the unbound 3TIH crystal)* while it may leave it exposed to thioredoxin after CD4 binding as observed in 4R4N chain B in table 2. This data may well explain Azimi’s finding that cleavage of the gp120 disulfide bond by thioredoxin is ehananced by the binding of gp120 to soluble CD4 [17]. Thiol/disulfide exchange affecting the Cys296-Cys331 disulfide bond at the base of the V3 loop is an acknowledged step in HIV-1 viral entry although the exact mechanism has yet to be elucidated [17, 19]. TTSB formation seems to protect the C296-C331 bond against thiolate attack in unliganded 3TIH whereas it tends to expose it to thiolate attack in 4R4N bound to CD4-mimetic. The transfer of energy between these allosteric disulfide bonds may be driven by factors not visible in our current crystal analysis. However the sequence of events in DSE distribution observed in the 3TIH and 4R4N crystals may pertain to a fine-tuning program for perfect timing of the Cys296-Cys331 cleavage to specifically occur after and not before CD4-binding.

Allosteric bonds are mechano-sensitive, reversible and capable of acting and reacting with mechanical forces [20] *(see tables 2,3 and 4)*. The TTSB intermediate formation could then apply the necessary timing program to control and orchestrate the flow of stored forces generated by these allosteric bonds to dictate which bond gets broken when.

**In summary** the TTSB formation represents an important intermediate structure to allow safe transfer of energy between these allosteric bridges involving Cys296, Cys331, Cys385 and Cys418, Cys378 and Cys445 for various possible functions such as shock absorber and/or more dynamic engagements like the bistable flip/switch phenomenon and/or V3 mechanics for viral entry. The TTSB connects to the LPCR motif via Cys418 and could act as a transmission for safe transfer of energy between the allosteric bonds to achieve safe movement within the molecule allowing the bistable switch phenomenon to occur without needing ATP expenditure while saving the integrity of structural disulfide bridging waiting till CD4 binding to allow proper timing for cleavage of the Cys296-Cys331 bridge at the base of the V3 loop to promote the fusion-active state conformational changes.

#### Literature review

Leonard et al. described the disulfide bond assignment in the gp120 molecule [21]. In the initial assignment of disulfide bridges in the gp120, there were a number of disulfide bridges that could not be resolved without persistent ambiguities, namely C348, C415, C388 and C355 in their tryptic maps. Eden Go et al. subsequently reported variety in the disulfide bond connectivity in the V1, V2 region of a gp140 molecule [22].

Site-directed mutagenesis studies proved that mutations at Cys 296, 331, 418 and 445 resulted in noninfectious mutants whose gp160 envelope precursor polypeptides were poorly cleaved [23] whereas mutations at Cys131, 196 resulted in noninfectious mutants that were defective for syncytium formation with a late block in life cycle (Tschachler et al. 1990) [24]. This same study also shows that viruses with mutations at Cys386 were still infective but defective for syncytium formation. Van Anken et al recently demonstrated that only 5 of the 10 conserved disulfide bridges in gp120 are essential for folding and 8 for function [25]. Of these, C296-C331 proved necessary for folding and function while C385-C418 mutants showed residual infectivity functions despite severe folding limitations; whereas mutations at C378-C445 bridge still allowed proper folding and infectivity. Sanders et al. subsequently demonstrated that C385A/C418A mutations (disrupting the C385-C418 disulfide bridge) can be overcome to resume infectivity in SupT1 cells and HeLA cells through evolution experiments by achieving a T415I mutation accompanied by substitutions of Valine in mutated positions 385 and 418, putting an emphasis on the beta-sheet strengthening for function [26]. The resilience of HIV is unsurpassed and continues to defy our current knowledge of chemistry. *“When you don’t know what you don’t know, you think you know”*: crystallization of this evolved gp120 mutant [26] is needed to help elucidate this phenomenon because the cysteine residues involved seem to have a significant importance in the life cycle of HIV-1.

While practical for the virus, the TTSB also presents a complex fragility for the gp120, which may explain why many antibodies that bind to the V3 loop –although strainspecific- (located between Cys296-Cys331 disulfide bridge involved in the TTSB) can easily prevent CD4-binding and neutralize HIV-1. Another complex fragility for the TTSB is the fact that this TTSB is attached, by way of Cys418, to the segment 416-427. As mentioned earlier this segment, which includes the LPCR motif, is involved in the bistable flip/switch phenomenon which appears to necessary for CD4-binding [5,6,7,8]. Moreover this segment [416-427] also contains the residues that are necessary for CD4-binding and CCR5-binding [9, 10, 27, 28].

The HIV-1 virus has many unusual properties that remain unexplained. Soluble gp120 by itself also has immunosuppressive effects [29]. Soluble gp120 inhibits T-cell-dependent B cell differentiation both at the inductive phase of T-cell activation and at the effector phase (Chirmule et al. 1990-1994) [30–37]. Soluble gp120 causes apoptosis in uninfected CD4+ cells. Humoral response to HIV gp120 can sensitize non-infected cells towards apoptosis and contributes to T cell decline upon administration of recombinant gp120 (Kang et al. 1998) [38]. Cerutti et al. subsequently demonstrated that the binding of gp120 to CD4 inhibits the thioredoxin-driven dimerization of CD4, which is a necessary step to allow the formation of functional MHCII-TCR-CD4 complex for antigen presentation [39]. Incidentally the binding of TCR to MHCII without the CD4-MHCII causes the T cell to undergo apoptosis. [36, 36].

While most of the mechanisms for these features of HIV-1 have yet to be elucidated, no immunomodulatory effect has ever been attributed to any sulfide bridge in the gp120. Gliotoxin and related epipolythiodioxopiperazines (ETP’s) have *in vitro* immunomodulating activities including apoptosis and inhibition of phagocytosis (Waring 1996) [40]. The immunoregulatory activities of gliotoxin and ETP’s have been attributed specifically to the sulfur bridges in these molecules (Mullbacher et al. 1986) [41]. The immunomodulating effects of these agents are prevented *in vitro* in the presence of reducing agents.

Given the observation of various TTSB configurations as Cys331-Cys385 and Cys331-Cys418, the triangular-shaped TTSB observed in the 3TIH crystal chain C clearly represents an important intermediate and it suggests a dynamic motion between the Cys385 and the Cys418 TTSB configurations. Which direction does it go ? does it go from 385 to 418 formation or vice versa ? does it go back and forth ? The direction and purpose of this dynamic motion have yet to be elucidated. However based on the DSE changes engendered by the different conformations as illustrated in table 2, TTSB intermediate formation proves to be important for safe energy transfer between the allosteric bonds. This safe transfer of energy between the allosteric bonds could allow the gp120 molecule to achieve various functions fine-tuning the steps for CD4-binding and viral entry mechanisms.

#### TTSB inhibitors

TTSB formation as presented in this study seems to be an important intermediate in gp120 dynamics be it for structure stabilization, CD4-binding, and/or viral entry mechanisms. Hence inhibition of TTSB formation may provide new therapeutic strategies for prophylaxis and treatment of HIV infection. Given the fact that these allosteric disulfide bonds are well conserved in all clades of HIV, TTSB inhibitors may achieve high genetic barriers for less risk of viral resistance in therapy against HIV with inter- and intra-clade coverage. Other viruses that use TTSB formation in cell entry mechanisms would likely be sensitive to TTSB inhibitors. As such, TTSB inhibitors could do for virology what penicillin does for bacteriology: broad-spectrum coverage. In addition, any toxin or venom which uses TTSB formation in their mechanisms of action could immediately be neutralized by TTSB inhibitors. Moreover TTSB formation inhibitors may also allow testing of other viruses and toxins that may involve TTSB formation in their mechanisms for cell entry and other functions before we could even capture such phenomena on crystallography.

### Conclusion

Since tetrasulfide bridges are not well studied in biological systems, it is not possible to enumerate the potential functions that could be attributed to these structures. While one could appreciate the potential chemical, biophysical and functional properties of TTSB formation even as a timing program, the dynamics of TTSB intermediates have yet to be elucidated. However the presence of such an extraordinary structure with potential to safely transfer energy between the allosteric bonds at the base of the V3 and V4 loops and at the base of the segment assigned to the bistable switch phenomenon does point to some intriguing insights in the multi-functional design and mechanics of the gp120 molecule…

## Acknowledgements

I thank God for guiding my steps in this endeavor in the art of caring. I also thank my family, friends and colleagues who do not always understand but often appreciate and share the enthusiasm in the thrill of discovery…

